# A Draft Genome for *Hirudo verbana*, the Medicinal Leech

**DOI:** 10.1101/2020.12.08.416024

**Authors:** Riley T. Paulsen, Diing D.M. Agany, Jason Petersen, Christel M. Davis, Erik A. Ehli, Etienne Gnimpieba, Brian D. Burrell

## Abstract

The medicinal leech, *Hirudo verbana*, is a powerful model organism for investigating fundamental neurobehavioral processes. The well-documented arrangement and properties of *H. verbana*’s nervous system allows changes at the level of specific neurons or synapses to be linked to physiological and behavioral phenomena. Juxtaposed to the extensive knowledge of *H. verbana’s* nervous system is a limited, but recently expanding, portfolio of molecular and multi-omics tools. Together, the advancement of genetic databases for *H. verbana* will complement existing pharmacological and electrophysiological data by affording targeted manipulation and analysis of gene expression in neural pathways of interest. Here, we present the first draft genome assembly for *H. verbana*, which is approximately 250 Mbp in size and consists of 61,282 contigs. Whole genome sequencing was conducted using an Illumina sequencing platform followed by genome assembly with CLC-Bio Genomics Workbench and subsequent functional annotation. Ultimately, the diversity of organisms for which we have genomic information should parallel the availability of next generation sequencing technologies to widen the comparative approach to understand the involvement and discovery of genes in evolutionarily conserved processes. Results of this work hope to facilitate comparative studies with *H*. *verbana* and provide the foundation for future, more complete, genome assemblies of the leech.

## Introduction

The medicinal leech (*Hirudo verbana*) has been repurposed from an ancient bloodletting instrument [1] to a widely utilized invertebrate model system in biomedical research [2]. The well-documented and accessible central nervous system of the leech allows for precise selection of neurons for electrophysiological studies based on their characteristic morphologies, positioning, and biophysical properties [3, 4]. Fundamental discoveries have been made using *Hirudo* in a variety of disciplines that include central pattern generators, behavioral choice, learning and memory, synaptic signaling, neuroethology, neuro-injury and repair, and neurodevelopment [5–9]. Extensive research has also been devoted to examining the proteins secreted during leech hematophagy, which has longstanding applications in inflammation and coagulation [10]. Genomic insights into the medicinal leech will facilitate a more comprehensive approach into the evolutionary conservation of genes involved in the mechanistic processes that the medicinal leech has been used to help elucidate.

Despite these well-documented advantages of the medicinal leech for addressing various research questions, the leech lacks the molecular and genetic tools in comparison to alternative model organisms [11, 12]. For example, *Caenorhabditis elegans* and *Drosophila melanogaster* have extensive resources for targeted genome engineering in addition to optogenetic tools for electrophysiology and behavior manipulation [13–16]. Improving the genomic resources of organisms like the medicinal leech will promote more inclusive comparative genomics approaches to identifying conserved structural and functional gene signatures involved in human health and disease. Moreover, for many years, the medicinal leech community had been inadvertently aggregating four species of medicinal leeches: *H*. *medicinalis*, *H*. *verbana*, *H. orientalis*, and *H*. *troctina* [17–20]. This misunderstanding regarding the taxonomic classification of these leech subspecies has led to some confusion surrounding appropriate cataloging of preliminary leech omics databases [21, 22]. Finally, in spite of the advancements in sequencing technology, most of the existing sequence repositories for the medicinal leeches have been comprised of expressed sequence tag [23] and transcriptomic databases [24–26], with recent leech genomics work centering around *H*. *medicinalis* despite the prominence of *H*. *verbana* in neuroscience research [27, 28]. Moreover, other leech genomes are also becoming available, including that for the Asian Buffalo leech, *Hirudinaria manillensis* [29]. This work presents the first draft genome for *H*. *verbana,* which consists of 250 Mbp, 61,282 contigs, an N50 of 8,638 bp, and a GC content of 38%. This draft genome, in addition to the growing transcriptomic resources for *H*. *verbana* [24, 26, 30] and the aforementioned genomic databases for the closely related leech species, will help accelerate studies seeking to link the molecular basis of previous and ongoing functional studies utilizing medicinal leeches.

## Materials and Methods

### Tissue collection and DNA extraction

High molecular weight genomic DNA was isolated from muscle of three specimens of *H. verbana* (obtained from Niagara Leeches, Cheyenne, WY) in separate preparations using the QIAamp DNA Mini Kit Cat. No. 51304 (Qiagen; Hilden, Germany). In accordance with the protocol, 25 mg of dissected muscle tissue from the body wall was used as input for each of the DNA extractions. The DNA from the three different isolations was pooled, concentrated with Ampure XP beads (Beckman Coulter; Brea, CA) and 500 ng (10 ng/µL) was utilized for sequencing library preparation.

### Library preparation

Sequencing libraries were prepared using the TruSeq Synthetic Long-Read DNA Library Prep Kit (Illumina, Inc; San Diego CA). Three sequencing libraries were prepared by loading the sample into a Covaris g-TUBE (part no. 520079), and centrifuging twice at 4200 x g for 1 minute. Briefly, the DNA was fragmented to approximately 8-10 kb and ligated with adapters, which mark the end of contigs during data analysis. Following a dilution to limit the number of DNA molecules in each well of a 384-well plate, long-range PCR was performed to enrich for DNA fragments with appropriate adapters. The DNA in each well was treated with the Nextera transposome, which fragments and simultaneously adds adapters to DNA. Indexing-PCR was used to barcode the DNA in each well of the 384-well plate. The resulting products were pooled and bead size-selection was performed. The average size of the final libraries was ∼725 bp as measured with a High Sensitivity DNA chip on a 2100 Bioanalyzer (Agilent; Santa Clara, CA). The concentration of each library was determined by quantitative PCR (qPCR) via the KAPA Library Quantification Kit for Next Generation Sequencing (KAPA Biosystems; Woburn, MA).

### Whole genome sequencing

Libraries were normalized to 2 nmol/L in 10 mM Tris-Cl, pH 8.5 with 0.1% Tween 20. Prior to cluster amplification, the libraries were denatured with 0.05 N NaOH and diluted to 20 pmol/L. Paired-end cluster generation of denatured templates was performed according to the manufacturer’s instructions (Illumina, San Diego, CA) utilizing the HiSeq Rapid PE Cluster Kit v2 chemistry and flow cells. Libraries were optimally clustered at 11 pmol/L with a 1% PhiX spike-in. Sequencing-by-synthesis was performed on a HiSeq 2500 utilizing v2 chemistry with paired-end 101 bp reads and an 8 bp index read.

### Long read and genome assembly

A total of 1,862,297,140 bp of 2 x 101 bp reads were obtained from three flow cells. Sequence read data were processed and converted to short-read FASTQ format by Illumina BaseSpace analysis software (v2.0.13). The short reads from each plate were individually processed in three runs to construct primary contigs using the TruSPAdes assembly software (v1.1.0) [31], which were merged into one assembly using CLC-Bio Genomics Workbench *De Novo* Assembly (Qiagen, v11.0.1) with default parameters. Thorough quality control was performed on the raw short read data using FastQC (http://www.bioinformatics.babraham.ac.uk/projects/fastqc/) to assess the Phred score, presence of repeat reads, non-nucleotide content, GC content, and duplicated read contents. The quality of the primary contigs assembled by the TruSPAdes algorithm were assessed by Quast [32].

### Repeat sequence and BUSCO annotation

Repeatmasker [33] was used to annotate repeating sequences and transposable elements in the *H. verbana* genome assembly. Repeatmasker was configured with the pooled databases RepBase-2017 [34] and Dfam-2017 consensus databases [35], and Tandem Repeats Finder, ran with RMBlast (v2.2.27) search engine. Additionally, the completeness and quality of the draft genome was evaluated using a BUSCO (Benchmarking Universal Single-Copy Orthologs) assessment that matched our newly assembled sequences to the metazoan OrthoDB v9.1 [36–38]. BUSCO.v3 was configured using AUGUSTUS gene predictor [39], HMMER [40, 41], and NCBI-BLAST+ [42].

### Functional annotation and orthologous analysis

In order to elucidate functional annotation and gene ontology annotation, NCBI Blast+, UniprotKB [43], and the Blast2GO software suite [44] integrated with InterProScan [45] were implemented. Furthermore, through locally constructed databases in CLC-Bio Genomics Workbench, we utilized NCBI Blastn [46] on our genome against closely related databases for the following closely related polychaete annelids: *H. medicinalis* (GenBank: EY478949-EY505781), *Helobdella robusta* (GCA_000326865.1), and *Capitella teleta* (GCA_000328365.1).

### Gene prediction and macro-synteny analysis

Gene predictions were performed using the MAKER2 [47] genome annotation pipeline with SNAP [48] against the nematode *Caenorhabditis elegans* and two other annelids: *C. teleta*, and *H. robusta*. The *ab-initio* gene predictor, SNAP, was trained three times to improve performance. The *H*. *verbana* draft genome assembly was aligned to the *C. elegans* genome and scaffolds of *H. robusta*. Circos [49] was used to generate the circular genome alignment figures to analyze the anchoring of the top 600 *H*. *verbana* contigs onto the six chromosomes of *C. elegans* [50]. Additonally, the draft genome top 600 contigs were mapped and anchored to the 1,991 scaffolds of the *H. robusta* genome.

### Phylogenetic reconstruction

OrthoFinder2 [51] was utilized for comparative genomics between our draft genome for *H*. *verbana*, and the protein sequence databases for six other organisms: *H. medicinalis*, *H. robusta*, *C. teleta*, *C. elegans* (GCA_000002985.3), and the chordates *Mus musculus* (GCA_000001635.8) and *Homo sapiens* (GCA_000001405.27). Orthofinder was configured with the DIAMOND search engine [52], MCL clustering algorithm [53], and FastTree [54] to construct the rooted phylogenetic tree which was visualized with Phylo.io [55].

### Species Identification

The presence of mitochondrial contigs was assessed by comparing the draft mitochondrial genomes for *H. medicinalis* and *H. verbana* [56] to our draft genome. NCBI Blastn [46] was used on our draft genome against the mitochondrial DNA sequences downloaded from NCBI GenBank. The phylogenetic relationships between the mitochondrial contigs present in our draft genome, *H*. *medicinalis*, *H. verbana*, as well as other Hirudinea, *Whitmania pigra* [57] and *Hirudo nipponia* [58] via their mitochondrial genomes was also analyzed. The evolutionary analysis was performed in MEGA X (v11.0) [59, 60] using Multiple Sequence Comparison by Log-Expectation (MUSCLE) [61], and the phylogenetic tree was constructed using the Maximum Likelihood method with the Tamura-Nei model [62] and node confidence was assessed with a 500 bootstrap trials.

## Results and Discussion

Next-generation sequencing leveraging an Illumina HiSeq-2500 platform was employed to construct the first draft genome for *H*. *verbana*, the medicinal leech. A total of 188 Gbp were generated that encompassed 1,862,297,140 bp of 101bp x 2 paired short reads (Table 1A). Quality control of the raw short reads performed by FastQC and MultiQC [63] revealed that the data had an average phred score >30 (S1 Appendix). The short reads were barcode assembled individually for each plate using TruSPAdes assembler software into TruSeq synthetic long reads. The long reads (Table 1B) were 190,514 bp, 198,741 bp, and 193,658 bp for each plate, respectively, and had an average N50 of 7623 bp. Together, these synthetic long reads consisted of 582,913 sequences, 3,429,493,670 bp, and had an estimated coverage of 6.9X. Quast assessment of both the synthetic long reads and final assembly maintained a phred score of 30. The draft genome was constructed from the TruSeq synthetic long reads using CLC-Bio Genomics Workbench to produce the draft genome assembly for *H*. *verbana*. Prior to arriving at the final assembly that used a combination of TruSPAdes synthetic long reads and CLC-Bio Genomics Workbench, we performed a thorough assessment of multiple assemblers using long reads formed both by Illumina BaseSpace analysis software and TruSPAdes in conjunction with SOAPdenovo2 [64], Megahit [65], Spades [66], Ray [67], and Velvet [68]. Ultimately, we are reporting the TruSPAdes and CLC-Bio assembly approach because it had the most coverage and performed best under downstream assessment described below.

**Table 1:** Statistics of the de novo draft genome assembly for Hirudo verbana. (A) Whole genome sequencing reads obtained from Illumina HiSeq for H. verbana de novo draft genome assembly (B) Statistics for barcode-assembled synthetic long reads generated using TruSPAdes (C) Summary of assembly statistics of the draft genome for H. verbana.

|  |  |  |  |
| --- | --- | --- | --- |
| (A) Genome sequencing reads obtained from Illumina HiSeq via Basespace |  |  |  |
|  | Reads (Single, 101bp) | Reads (Pair, 101bp x 2) | Bases (Pair) |
| Plate 1 | 266,688,108 | 533,376,216 | 53,870,997,816 |
| Plate 2 | 328,593,254 | 657,186,508 | 66,375,837,308 |
| Plate 3 | 335,867,208 | 671,734,416 | 67,845,176,016 |
| Total reads | 931,148,570 | 1,862,297,140 | 188,092,011,140 |
| (B) TruSPAdes Truseq Synthetic Long-Read Assembly Statistics |  |  |  |
|  | Assembled reads | Total bases | N50 |
| Plate 1 | 190,514 | 1,117,298,449 | 7,612 |
| Plate 2 | 198,741 | 1,172,622,590 | 7,634 |
| Plate 3 | 193,658 | 1,139,572,631 | 7,624 |
| (C) Summary details of *Hirudo verbana* draft genome assembly |  |  |  |
| Estimated genome size | 250 Mb |  |  |
| Total assembly length | 250 Mb |  |  |
| Total sequences | 582,913 |  |  |
| Short read coverage | 627X |  |  |
| Long read coverage | 6.9X |  |  |
| Contigs | 61,282 |  |  |
| N75 | 4807 bp |  |  |
| N50 | 8638 bp |  |  |
| N25 | 14800 bp |  |  |
| Minimum contig | 200 bp |  |  |
| Maximum contig | 154993 bp |  |  |
| Average | 4084 bp |  |  |
| Nucleotide frequency |  |  |  |
| Adenine (A) | 30.90% |  |  |
| Cytosine (C) | 19.10% |  |  |
| Guanine (G) | 19.00% |  |  |
| Thymine (T) | 31.00% |  |  |

The final draft genome assembly presented here is 250,270,938 bp in size, which is comparable to the genome assembly statistics for other annelids *C. teleta* (240 Mbp) and *H*. *robusta* (310 Mbp). The genome assembly consists of 61,282 contigs that have a minimum length of 200 bp, a maximum length of 154,993 bp, and an N50 of 8,638 bp (Table 1C). N50 is important in an assembly assessment as it measures contigs lengths distribution and informs on the internal parameter of the assembly. However, it does not inform or guarantee the goodness/completeness of an assembly [69]. For example, filtering out of a shortest contigs or the scaffolding that links contigs could increase the N50 without improving the assembly completeness. Also, when analyzing the N50, a comparison should be made between draft genomes of a similar length, or it will be misleading [70]. In this assembly, we picked the assembly run with 200 bp parameter setting as the shortest contig cutoff. While assemblies were tested with higher cutoffs, specifically 500 bp and 1000 bp, the chosen setting produced the best outcome in in terms of balance of genome size coverage and N50 criteria. A condensed summary comparing our draft genome to the 10 available annelid genomes is also provided in S1 Table.[27-29, 71-76] For preliminary validation of the quality of the draft genome assembly, the 61,282 contigs were mapped back to the raw short reads using the mapping module in CLC-Bio Genomics Workbench. The result demonstrated that 86.72% of the assembly mapped back onto the short reads, leaving 13.28% unmapped. Among the contigs that were reported to map back, 85.77% had identical base pair matching.

Next, the repeating segments and transposable elements of the draft genome were annotated using RepeatMasker. An estimated 6.67% of the genome assembly (16,685,142 bp of the total 250 Mbp) was determined to be repetitive or transposable elements, with a majority consisting of simple repeats, interspersed repeats, and low complexity repeats (S2 Table). Moreover, the completeness of the draft genome was assessed with a BUSCO analysis for the presence of metazoan-specific orthologues. The metazoan BUSCO that we implemented consisted of 978 genes, and our assembly returned 809 (82.70%) as complete, 533 (54.50%) as complete and single-copy, 276 (28.20%) complete and duplicated, 70 (7.20%) fragmented, and 99 (10.10%) as missing (Table 2A). Overall, our genome has a completeness score of 89.9% (82.70% complete + 7.20% fragmented).

**Table 2.**
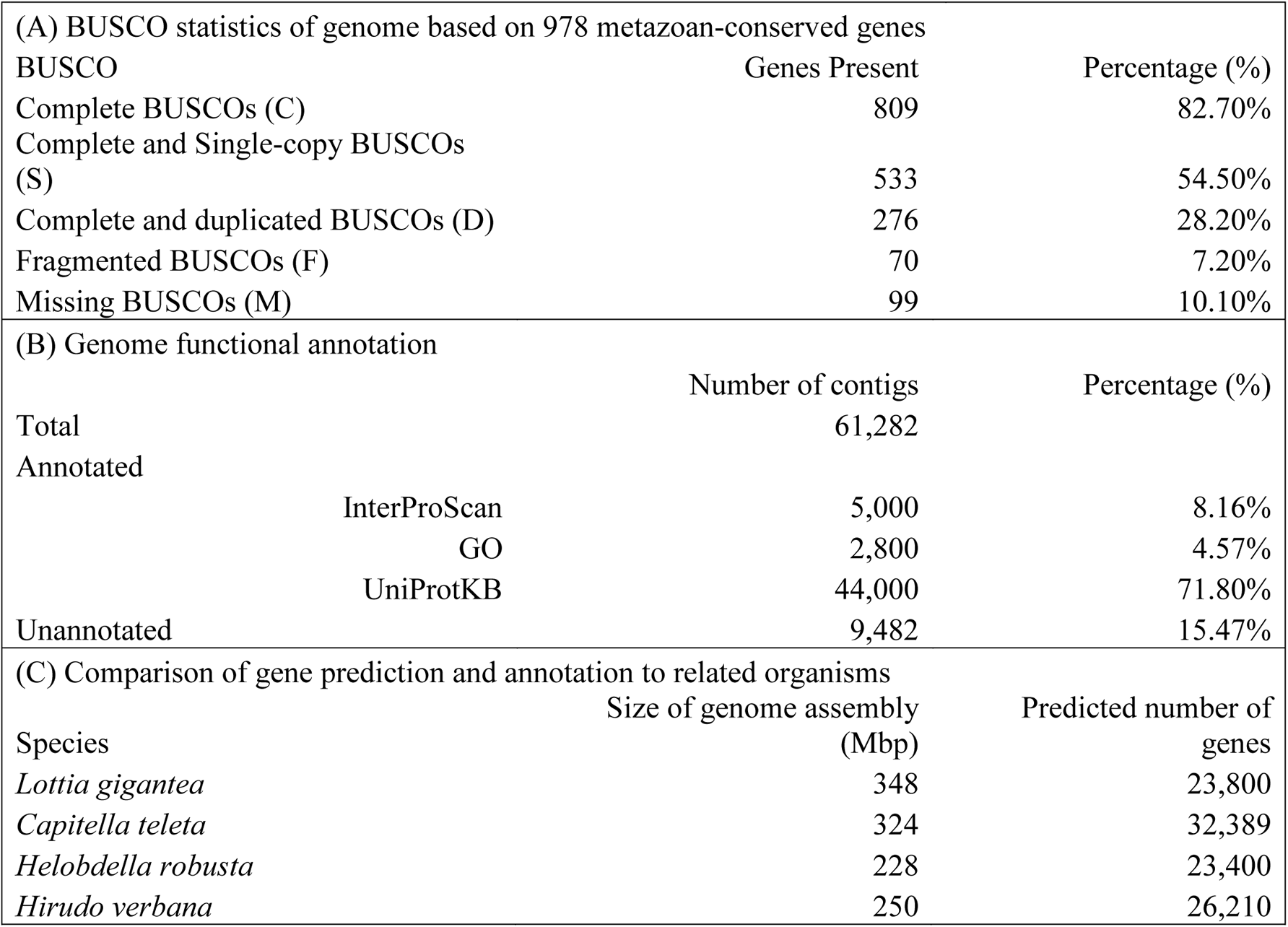
Analysis of the completeness and orthology of the H. verbana genome. (A) BUSCO statistics assessing the completeness of the H. verbana draft genome based on 978 metazoan-conserved genes (B) Summary of structural functional annotation for H. verbana draft genome (C) Comparison of H. verbana draft genome size and predicted number of genes to 3 closely related spiralian genomes (two annelids and one mollusc).

|  |  |  |
| --- | --- | --- |
| (A) BUSCO statistics of genome based on 978 metazoan-conserved genes |  |  |
| BUSCO | Genes Present | Percentage (%) |
| Complete BUSCOs (C) | 809 | 82.70% |
| Complete and Single-copy BUSCOs (S) | 533 | 54.50% |
| Complete and duplicated BUSCOs (D) | 276 | 28.20% |
| Fragmented BUSCOs (F) | 70 | 7.20% |
| Missing BUSCOs (M) | 99 | 10.10% |
| (B) Genome functional annotation |  |  |
|  | Number of contigs | Percentage (%) |
| Total | 61,282 |  |
| Annotated |  |  |
| InterProScan | 5,000 | 8.16% |
| GO | 2,800 | 4.57% |
| UniProtKB | 44,000 | 71.80% |
| Unannotated | 9,482 | 15.47% |
| (C) Comparison of gene prediction and annotation to related organisms |  |  |
| Species | Size of genome assembly (Mbp) | Predicted number of genes |
| *Lottia gigantea* | 348 | 23,800 |
| *Capitella teleta* | 324 | 32,389 |
| *Helobdella robusta* | 228 | 23,400 |
| *Hirudo verbana* | 250 | 26,210 |

The sequence homology and similarity of the draft genome for *H*. *verbana* was assessed by NCBI BLAST+ against the genomic and transcriptomic sequences available for *H*. *medicinalis*, *H*. *robusta*, and *C*. *teleta*. Approximately 94% of the draft genome sequences had an identity match within the queried databases, 5.5% exhibited at least 70% similarity, and the remaining 0.5% was unidentified. Functional annotation was assessed using fast-BlastX [77] against the Animalia NCBI Refseq database in Blast2GO [78]. From total assembly, 1,178 contigs returned significant blast hits at an e-value threshold of 10^-10^. The top 20 gene ontology (GO) terms [79] for each classification – biological process (BP), molecular function (MF), and cellular component (CC) – at GO level 5 are displayed in Fig 1. Moreover, the draft genome was aligned to the genome of *C*. *elegans*. The draft genome contigs for *H*. *verbana* were mapped and anchored to the 6 chromosomes of *C*. *elegans* (Fig 2A) and to the *H. robusta* scaffolds at position 15000-17000 for contig *hv*7 on scaffold *he49* (Fig 2B). A majority of the mapped sequences achieved better fit onto chromosomes 1 and X of the *C*. *elegans* genome (S3 Table).

**Figure 1.**
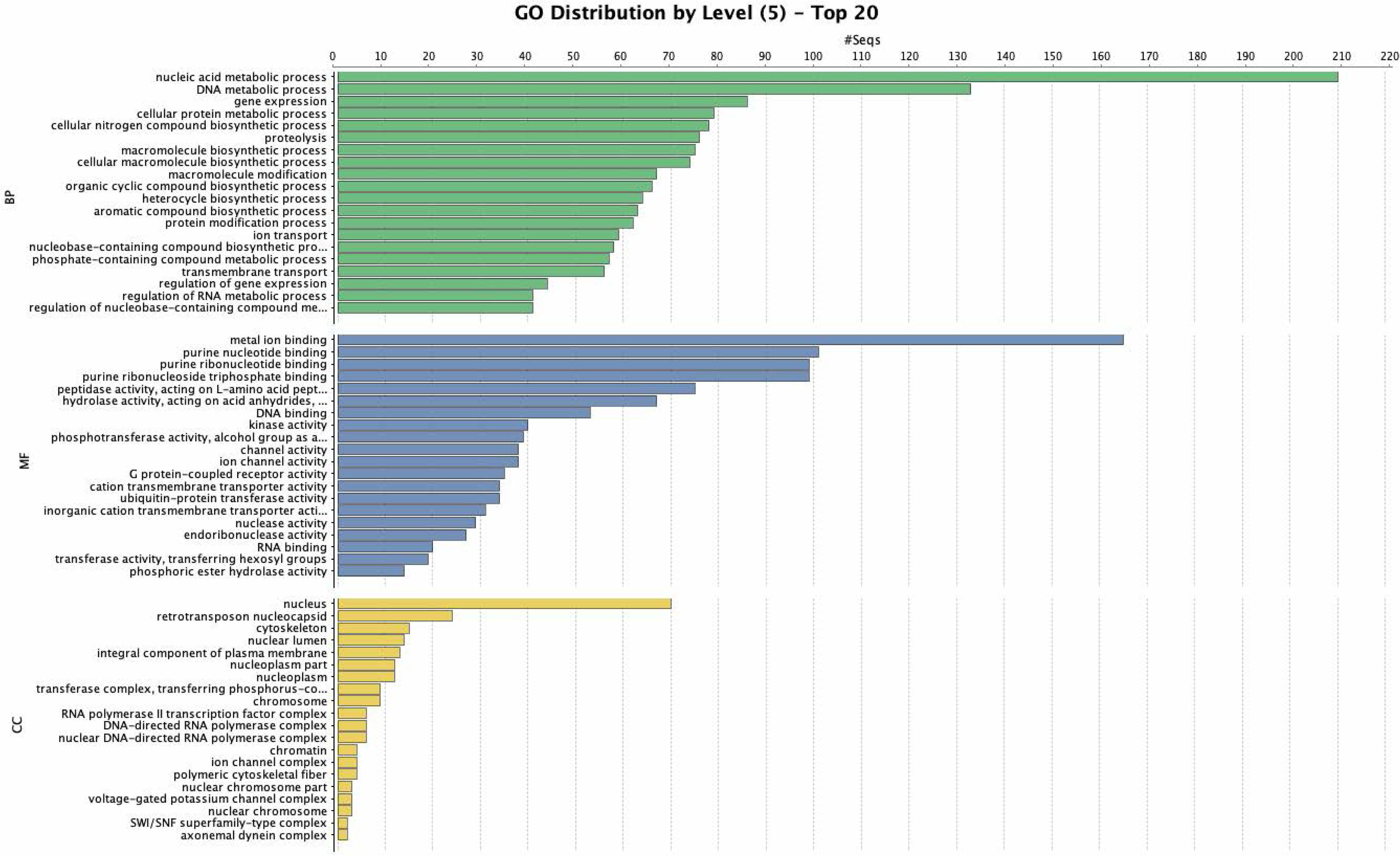
*H. verbana* draft genome gene ontology distribution. Displayed are the number of sequences present corresponding to the top 20 most frequent annotations for each classification. Gene ontologies shown are grouped into the major categories of: biological process (BP, green), molecular function (MF, blue), and cellular component (CC, yellow) at a gene ontology level of 5.

**Figure 2.**
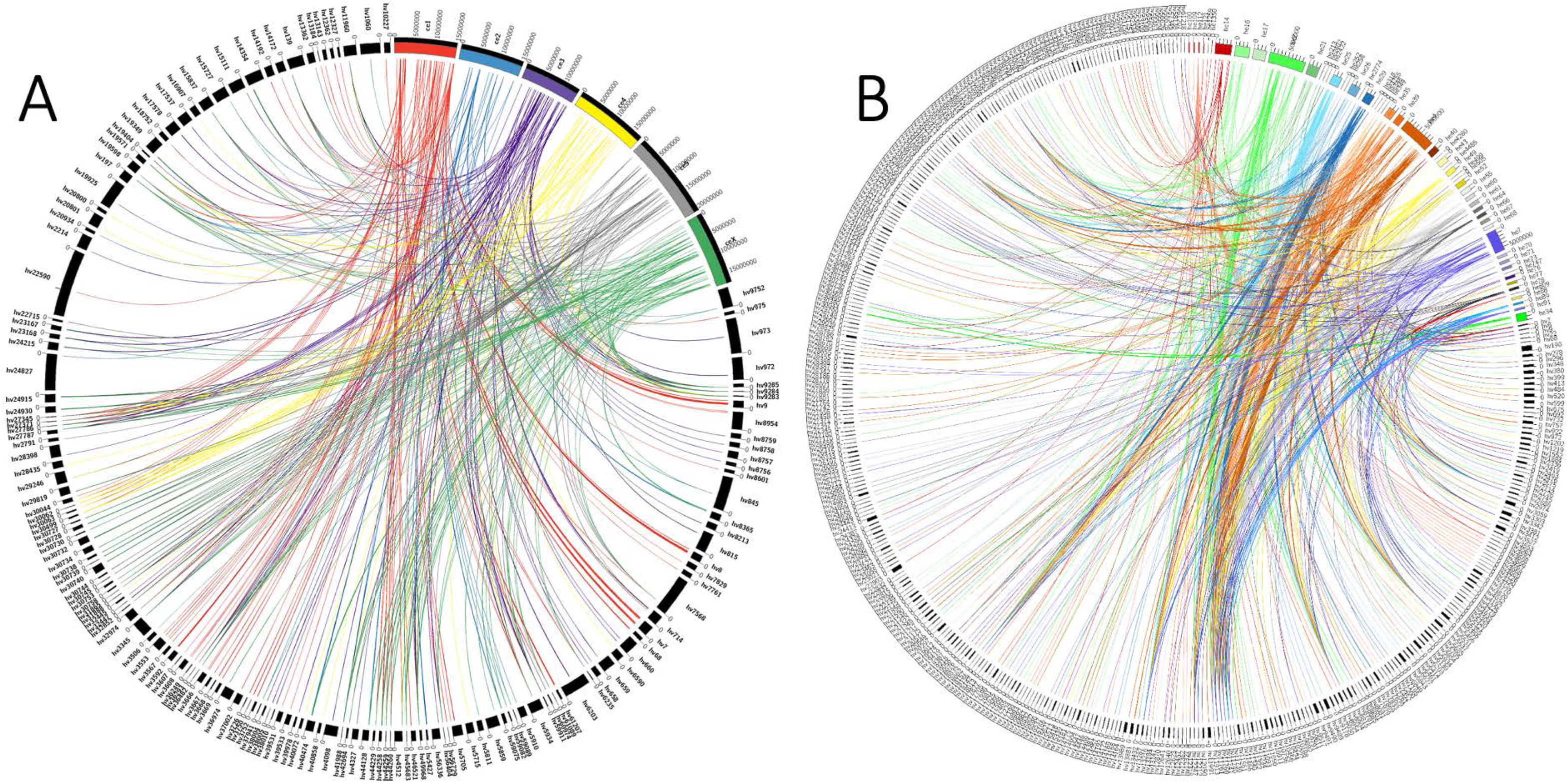
Alignment of *H. verbana* draft genome to *C. elegans* and *H. robusta*. (A) Alignment (>100 bp) of top 600 *H*. *verbana* draft genome contigs to the chromosomes (I-X) of *C*. *elegans*. (B) Alignment (>100 bp) of the top 600 *H. verbana* draft genome contigs to the scaffolds of the *H. robusta* genome assembly. *H. verbana* contigs are in black and abbreviated as “Hv,” the *C. elegans* chromosomes are abbreviated “ce,” and the *H. robusta* scaffolds are abbreviated “he.” Both the corresponding chromosomes and scaffolds are colored, and the projections are anchored to different contigs from the draft genome.

Using OrthoFinder, orthologues were generated from gene families of our draft genome for *H*. *verbana*, *H*. *medicinalis*, *H*. *robusta*, C*. teleta*, *C. elegans*, *M. musculus*, and *H. sapiens*. The phylogenetic tree was reconstructed using OrthoFinder after it identified the highest similarity content between the draft genome and the reference organisms. The phylogenetic tree (Fig 3A) appropriately placed *H*. *verbana* adjacent to its closest relative, *H*. *medicinalis*, demonstrating their last known divergence in genus *Hirudo*. Predictive protein-coding gene sequences were identified based on conserved protein signatures and domains with UniprotKB, Blast2Go, and InterProScan. A total of 84.53% of the contigs were annotated for a protein-coding function, of these, 8.16% were identified by InterProScan, 4.57% by Blast2GO, and 71.80% by UniProtKB (Table 2B). Lastly, two-pass annotation in MAKER predicted 26,210 protein-coding genes in the draft genome, which is similar to the first reports of draft genomes for fellow lophotrochozoans *C*. *teleta*, *H*. *robusta*, and *L. gigantea,* a gastropod mollusc (Table 2C).

**Figure 3.**
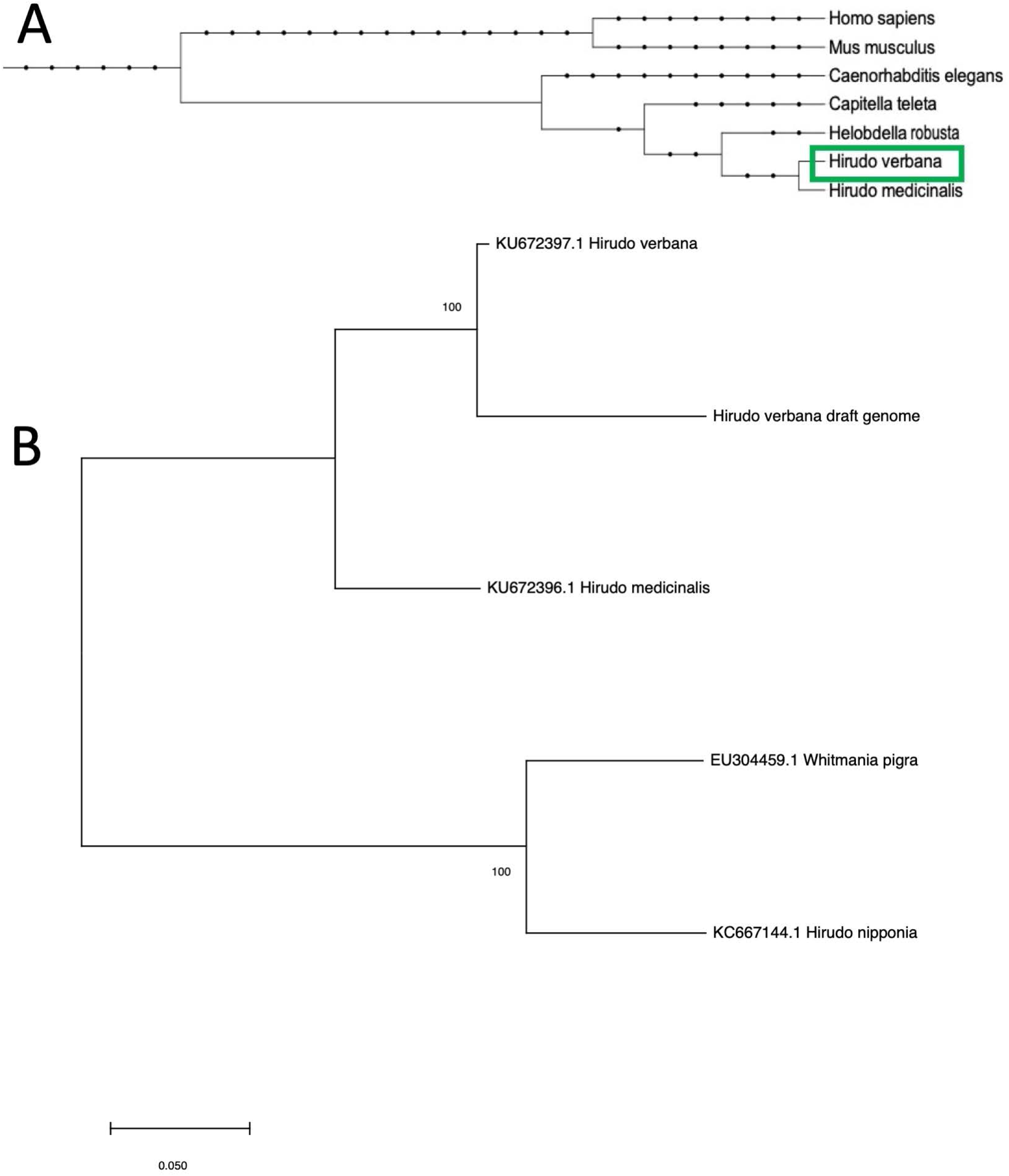
Genomic and mitochondrial phylogenetic relationships. (A) Reconstructed phylogenic tree based on orthologous gene families. (B) Phylogeny of our draft genome in relation to 4 leech species by comparing the presence and similarity of mitochondrial contigs through the Maximum Likelihood method via the Tamura 3-parameter model. NCBI accession numbers are provided next to each given taxa name.

For further confidence that the leech used in this study were *H*. *verbana*, we searched our draft genome for mitochondrial contigs using the draft mitochondrial genomes for *H*. *medicinalis* and *H. verbana*. The results of the mitochondrial contig analysis suggest that the leeches used to collect genomic DNA in our study were of the species *H*. *verbana*, which is emphasized by the mitochondrial contigs found in our draft genome forming a clade with the *H*. *verbana* draft mitochondrial genome in the phylogenetic tree (Fig 3B). The results of the mitochondrial phylogenetic analysis, in addition to the characteristic coloration patterns observed on the dorsal side of the leeches, indicate that the leeches used in this study were *H*. *verbana*.

This study is the first to publish an annotated draft genome sequence for the medicinal leech, *H. verbana*. Overall, the genome assembly consists of 250 Mbp and 61,282 contigs, 84.53% of which have been predicted to contain a protein-coding function. The draft genome is also predicted to contain 26,210 protein-coding genes and a repetitive content of 6.67%. The raw short-read sequence data, synthetic long-reads, and assembled contigs for the present study have been deposited into NCBI accession number JAIQDV000000000. The draft genome assembly will assist in providing tools to understand the underlying molecular processes involved in ongoing studies in neurophysiology, developmental biology, and neuroethology that utilize *H*. *verbana* [80–83]. Whole-genome characterization for Hirudinae *H*. *medicinalis* [27, 28], *H*. *manillensis* [29] and *H*. *verbana* is expanding and will help distinguish and clarify distinct genetic undertones of these previously amalgamated species. Future efforts to better annotate and complete a genome for *H*. *verbana* will enable insight into genetic mechanisms of processes investigated with this model organism [83–85], and advance more robust cross-species validation of comparative principles in next-generation biomedical research techniques.

## Acknowledgements

This work was supported by a pilot grant from the University of South Dakota (USD) Center for Brain and Behavior Research (CBBRe), the USD Neuroscience, Nanotechnology, and Networks Program (DGE-1633213), and the National Science Foundation Graduate Research Fellowship (DGE-1545679). Authors acknowledge the use of computational resources of the USD HPC cluster and the Extreme Science and Engineering Discovery Environment (XSEDE), which is supported by the National Science Foundation grant number ACI-1548562.

## Supplementary Figure/Tables

**Supplementary Figure 1.** Quality Control from Sequencing to Final Assembly Stage. (A) Integrated quality assessment for short sequencing reads generated by FastQC and MultiQC (B) GC content for final assembly and (C) TruSPAdes synthetic long reads after barcode assembly.

**Supplementary Table 1.**
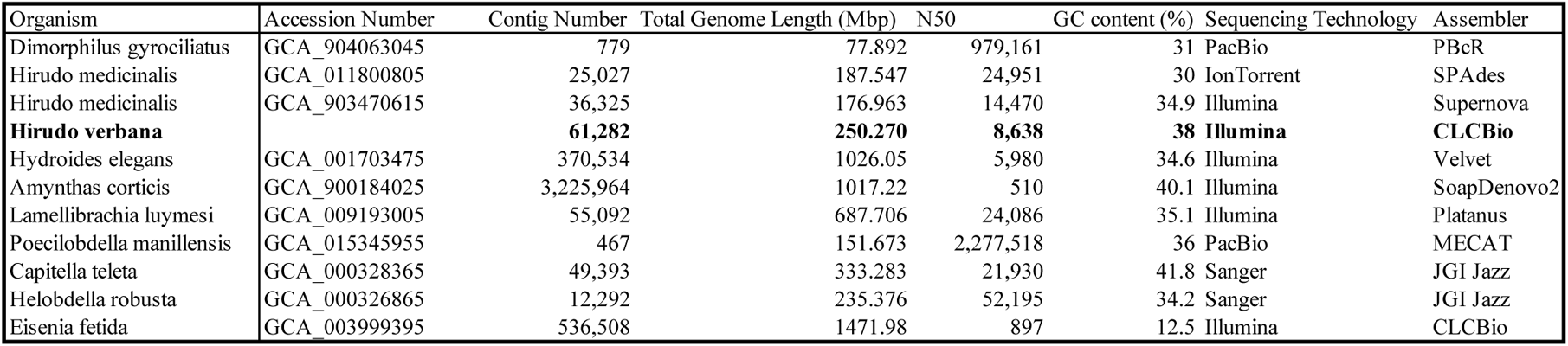
Annelid Genome Comparison. Comparison of select assembly statistics at the contig level (number of contigs, genome length, N50, and GC content) for annelid genomes. Accession numbers are provided, and the mode of sequencing technology and assembly software are also included for comparison.

**Supplementary Table 2.** Repeat sequence and transposable element composition in the draft genome of *H. verbana*. Identification of repetitive elements in the *H. verbana* draft genome using RepeatMasker.

| Element classification | No. of elements | Length (bp) | % of sequence |
| --- | --- | --- | --- |
| Total length of contigs | 61,282 | 250,270,938 |  |
| Repetitive sequences |  | 16,685,142 | 6.67% |
| Transposable elements |  |  |  |
| LTR | 327 | 107,597 | 0.04% |
| ERV_L | 36 | 8,360 | 0.00% |
| ERV_L-MaLRs | 31 | 10,905 | 0.00% |
| ERV_class I | 30 | 3,225 | 0.00% |
| ERV_class II | 2 | 110 | 0.00% |
| DNA | 870 | 132,616 | 0.05% |
| hAT-Charlie | 327 | 43,781 | 0.02% |
| TcMar-Tigger | 119 | 27085 | 0.01% |
| LINE | 3,202 | 622,089 | 0.25% |
| LINE1 | 112 | 35,062 | 0.01% |
| LINE2 | 1,378 | 316,959 | 0.13% |
| L3/CR1 | 1,411 | 237,912 | 0.10% |
| SINE | 286 | 36,605 | 0.01% |
| ALUs | 93 | 20,833 | 0.01% |
| MIRs | 77 | 8,440 | 0.00% |
| Unclassified | 76 | 5410 | 0.00% |
| Interspersed repeats |  | 904317 | 0.36% |
| Small RNA | 6 | 1261 | 0.00% |
| Satellites | 43 | 3896 | 0.00% |
| Simple repeats | 236935 | 15011804 | 6.00% |
| Low complexity | 15265 | 765808 | 0.31% |

**Supplementary Table 3.** Chromosome rearrangement of *C. elegans* onto *H. verbana* contigs. Mapping of *H. verbana* draft genome contigs onto the chromosome scaffolds of *C. elegans*.

| *C. elegans* chromosome | *Hirudo verbana* selected Contigs |
| --- | --- |
| **1** | hv3592, hv3669, hv3750, hv5705, hv61207, hv68, hv7, hv8, hv9, hv3668, hv3752 |
| **2** | hv5427, hv38001, hv18257, hv45683 |
| **3** | hv6203, hv972, hv973, hv32447, hv38005, hv39978, hv15111, hv17537, hv2214, hv24215, hv23168, hv17578, hv714, hv29246, hv38010, hv60974, hv8213, hv8757, hv8758, hv8756, hv8759, hv4098, hv27371, hv27345 |
| **4** | hv18257, hv714, hv29246, hv27371, hv27345 |
| **5** | hv815, hv14354, hv19349, hv24915, hv24930, hv30728, hv30732, hv30740, hv30739, hv30727, hv30744, hv30745, hv18257, hv22715, hv29819, hv2791, hv30738, hv30753, hv30730, hv30758, hv30734, hv42694 |
| **X** | hv14354, hv19349, hv24930, hv30728, hv30732, hv30740, hv30739, hv30727, hv18257, hv29819, hv42694, hv36974, hv5811, hv5859, hv30738, hv30753, hv30758, hv30734, hv30730, hv24215, hv23168, hv23167 |

